# Age-associated B cells are long-lasting effectors that restrain reactivation of latent γHV68

**DOI:** 10.1101/2021.12.29.474434

**Authors:** Isobel C. Mouat, Iryna Shanina, Marc S. Horwitz

## Abstract

Age-associated B cells (ABCs; CD19^+^CD11c^+^T-bet^+^) are a unique population that are increased in an array of viral infections, though their role during latent infection is largely unexplored. Here, we use murine gammaherpesvirus 68 (γHV68) to demonstrate that ABCs remain elevated long-term during latent infection and express IFNγ and TNF. Using a strain of γHV68 that is cleared following acute infection, we show that ABCs persist in the absence of latent virus, though their expression of IFNγ and TNF is decreased. With a fluorescent virus we demonstrate that ABCs are infected with γHV68 at similar rates to other previously activated B cells. We find that mice without ABCs display defects in anti-viral IgG2a/c antibodies and are more susceptible to γHV68 reactivation when challenged with heterologous infection. Together, these results indicate that ABCs are a persistent effector subset during latent viral infection that restrains γHV68 reactivation.

## Introduction

Humans are infected with an array of herpesviruses that persist within us throughout our lives and require continuous surveillance by the host immune system. Gammaherpesvirus-68 (γHV68), like other herpesviruses, deploys distinct transcriptional programs during lytic and latent infection, each of which requires a distinct immune response^1^. Age-associated B cells (ABCs) are a unique B cell population implicated in viral infection, autoimmunity, and aging^2–5^. ABCs (CD19^+^CD11c^+^T-bet^+^) are induced following γHV68 infection^6^, though their function throughout the course of infection is not clear. In this study we examine the response of and role for ABCs throughout γHV68 infection, from acute infection through long-term latency.

ABCs were identified in 2011 in the context of female aging and autoimmunity^2,3^ and have since been shown to be increased in an array of viral infections including lymphocytic choriomeningitis virus (LCMV), murine cytomegalovirus, γHV68, vaccinia, human immunodeficiency virus, rhinovirus, SARS-CoV2, and influenza^6–11^. ABCs are elevated in the spleen and circulation during active viral infections and persist primarily in the spleen during chronic infection or upon infection resolution^8,9^. ABCs display multiple functional capacities including the secretion of antibodies and anti-viral cytokines and activation of T cells^12,13^. T-bet expression in B cells is required for IgG2a/c class switching^14^ and lack of ABCs exacerbates LCMV chronic infection in mice^15^. We predict that ABCs could be playing a role throughout γHV68 infection due to their long-term persistence, activation of T cells, and continuous cytokine and antibody production.

Lytic γHV68 infection is primarily cleared by CD8^+^ T cells^16,17^, following which γHV68 persists in a latent state mainly in previously activated B cells^18^. During latency the immune response persists even though there is minimal production of viral genes and limited production of new virions^19^. Both humoral and cellular responses contribute to controlling γHV68 infection long-term^20,21^. Mice lacking either antibodies or T cells are both able to control γHV68 latency, though mice lacking both antibodies and T cells display elevated γHV68 reactivation^20,22^. IFNγ is critical for controlling γHV68 replication and reactivation from latency^23,24^.

B cells are also known to be important during latent γHV68 infection. B cells act as the primary latent viral reservoir, are required for movement from the lung to the spleen at the onset of latency, and mice deficient in B cells are unable to develop latency following intranasal infection^18,25–27^. B cells secrete anti-viral antibodies long term^28^, but also may inhibit reactivation via secretion of anti-viral cytokines or help to virus-specific T cells. The precise B cell subsets, mechanisms, and factors that facilitate the maintenance of latency and prevent reactivation are not fully understood.

The role of ABCs during acute and latent herpesvirus infection is unknown. Herein, using various tools, we investigate the relationship between γHV68 and ABCs throughout lytic and latent infection. We find that ABCs expand during acute γHV68 infection and persist in the spleen during latency. Well after the initial establishment of latency, ABCs continuously secrete anti-viral cytokines and antibodies and in the face of heterologous challenge, ABCs are required to restrain latent γHV68 reactivation. Thus, our findings highlight a novel role for ABCs as effector cells during γHV68 latency.

## Results

### ABCs are induced and maintained following γHV68 infection in a sex-biased manner

To examine the relative proportion of ABCs during acute and latent γHV68 infection, C57BL/6(J) mice were mock-infected with media (naïve) or infected with γHV68 for 6, 35, and 150 days, blood and spleen were collected and analyzed by flow cytometry to measure ABCs. A gating scheme, described in Figure 1A, was used to identify CD11c^+^T-bet^+^ as a proportion of previously activated B cells (CD19^+^IgD^-^). We observed the relative proportion of ABCs increased in circulation during acute γHV68 infection (6 days p.i.) compared to naïve mice (Figure 1B-C). Once latency was well established (day 35and 150 p.i.), the proportion of circulating ABCs decreased to pre-infection levels (Figure 1B-C). This indicates that ABCs circulate in response to the lytic γHV68 infection but do not remain elevated in circulation during latency. In the spleen, the relative proportion and total number of ABCs was increased during acute γHV68 infection (day 6 p.i.) relative to naïve mice and remained elevated during latent γHV68 infection at 35 days p.i. and 150 days p.i. (Figure 1D-E, Figure S1). These results support previous findings that the spleen is the major site for ABCs following the clearance of active viral infection^9^.

**Figure 1.**
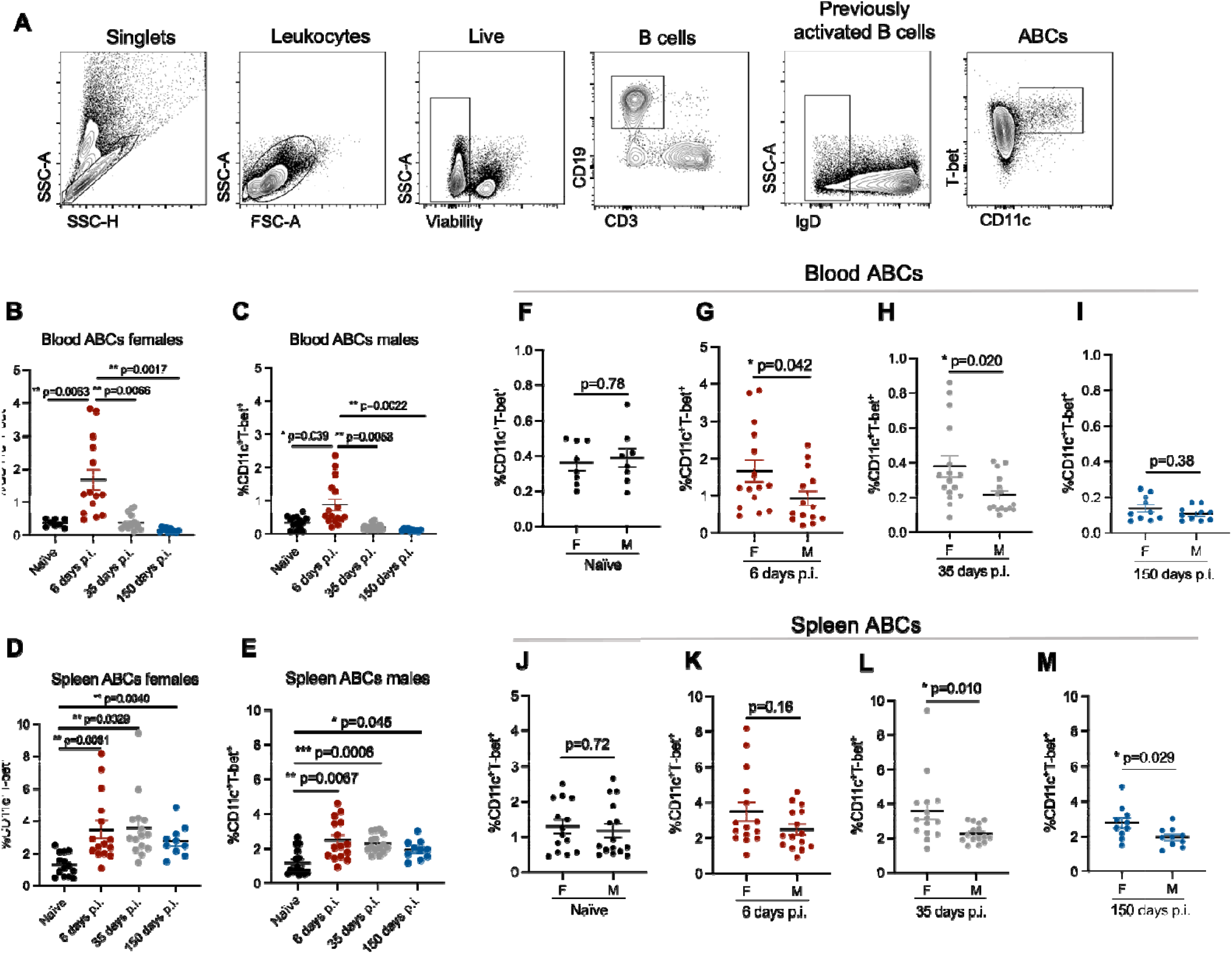
ABCs expand in the blood and spleen throughout γHV68 infection. Blood and spleen collected from C57BL/6(J) mice (6–8-week-old at infection) mock-infected with media (naïve, black circles) or infected i.p. with γHV68 for 6 (red circles), 35 (grey circles), or 150 (blue circles) days and processed for flow cytometry. (**A**) Representative gating strategy of ABCs. IgD, CD11c, and T-bet gating used fluorescence-minus-one (FMO) controls. (**B-E**) Proportion of ABCs (CD11c^+^T-bet^+^) of previously activated B cells (CD19^+^IgD^-^) in the blood and spleen of male and female mice. (**F-I**) Proportion of ABCs (CD11c^+^T-bet^+^) of previously activated B cells (CD19^+^IgD^-^) in the blood or spleen of male and female mice. Same data as presented in panels B-E. n=10-15 mice per group, data compiled from 4 experiments. Each data point represents an individual mouse. Data presented as mean ± SEM. Analyzed by (**B-E**) one-way ANOVA or (**F-M**) Mann-Whitney test. P-values indicated as asterixis as follows: *** p<0.001, ** p<0.01, * p<0.05.

Intriguingly, we observed that the proportion of ABCs is increased in females as compared to males in both the blood and spleen throughout infection (Figure 1F-M). Previously, we demonstrated that ABCs display a sex bias during experimental autoimmune encephalomyelitis and at 35 days post-γHV68 infection in the spleen^29^. Here, we have extended this finding and shown that ABCs display female sex bias in both the circulation and spleen during lytic and latent γHV68 infection. While ABCs increase in both females and males following γHV68 infection, there is a sex bias leading to greater expansion in females. These results demonstrate that ABCs are increased in circulation during acute γHV68 infection and continue to endure in the spleen long-term during latent infection.

### ABCs express anti-viral cytokines in response to latent γHV68 infection

To begin investigating the role of ABCs in the anti-viral response, we measured ABC expression of anti-viral cytokines interferon-γ(IFNγ) and tumor necrosis factor-α (TNF) in the spleen at 6, 35, and 150 days p.i.. We found that during acute γHV68 infection (6 days p.i.), roughly 40% of ABCs in the spleen expressed IFNγ and TNF, as compared to about 10% of ABCs in naïve mice (Figure 2A-B). During latent infection (35 days and 150 days p.i.), a significantly increased proportion of ABCs continued to express IFNγ and TNF compared to naïve mice, though the proportion of ABCs expressing IFNγ and TNF was decreased compared to acute infection (Figure 2A-B). Throughout the acute and latent infection, we observed a downregulation of IL-17A expression on ABCs (Figure 2C).

**Figure 2.**
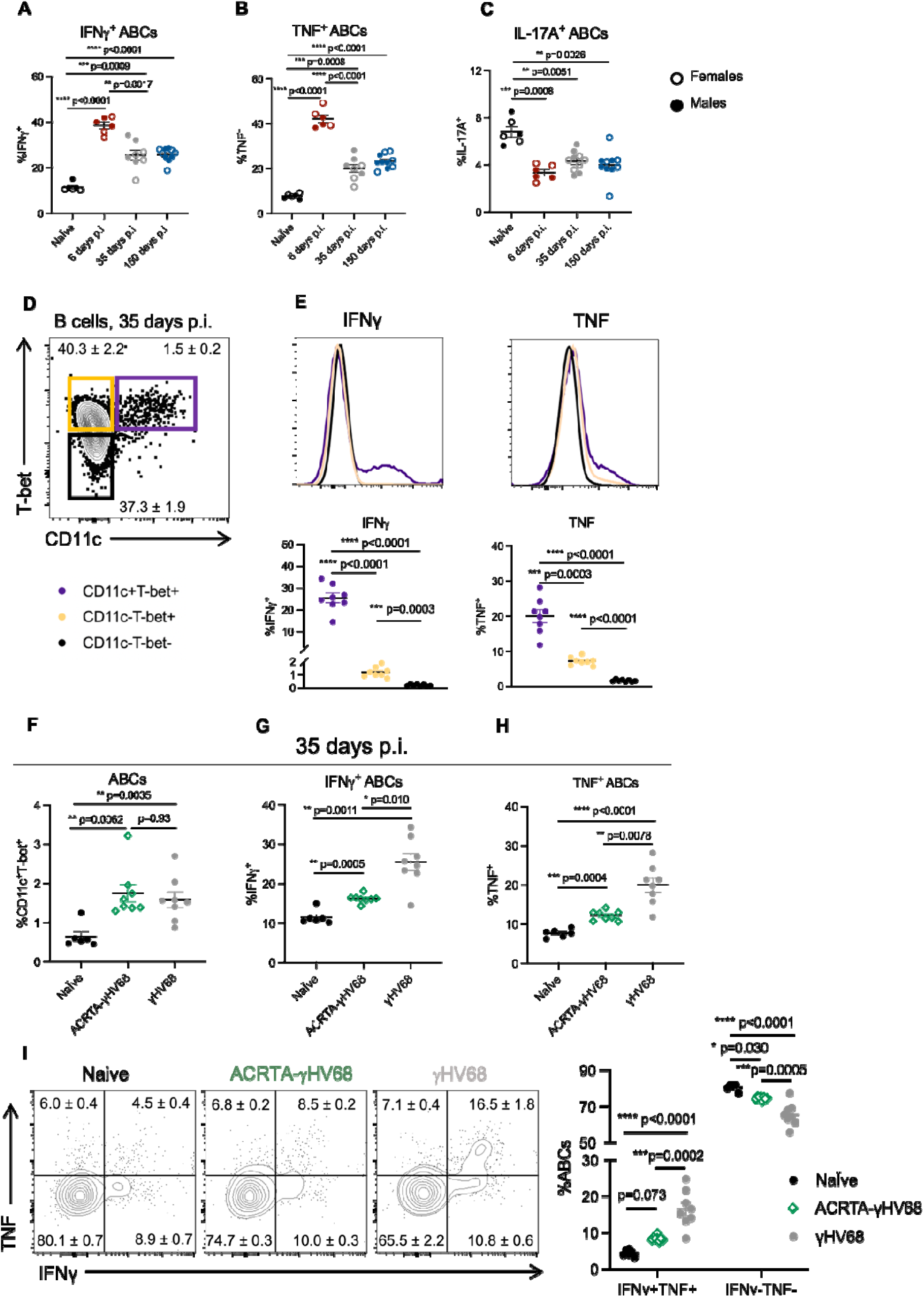
Cytokine production by ABCs in the spleen during γHV68 and ACRTA-γHV68 infection. **(A-C)** C57BL/6(J) mice (6–8-week-old at infection) mock-infected with media (naïve) or infected i.p. with γHV68 for 6, 35, or 150 days. Spleens collected and processed for flow cytometry. Proportion of ABCs in the spleen expressing (**A**) IFNγ, (**B**) TNF, or (**C**) IL-17A, with females presented as open circles and males as filled circles. **(D)** Gating of ABCs (CD11c^+^T-bet^+^) and non-ABC B cells (CD11c^-^T-bet^+^ and CD11c^-^T-bet^-^), previously gated on CD19^+^IgD^-^ cells in the spleen. **(E)** Representative histograms and dot plots of the proportion of ABCs and non-ABC B cells (CD11c^-^T-bet^+^ and CD11c^-^T-bet^-^) expressing IFNγ and TNF from mice infected with γHV68 for 35 days. (**F-H**) C57BL/6(J) mice mock-infected with media (Naïve, black circles) or infected i.p. with γHV68 (grey circles) or ACRTA-γHV68 (green diamonds) for 35 days and spleens processed for flow cytometry. (**F**) Proportion of ABCs (CD11c^+^T-bet^+^) of previously activated B cells (CD19^+^IgD^-^) in the spleen. Proportion of ABCs (CD19^+^CD11c^+^T-bet^+^) in the spleen expressing **(G)** IFNγ or **(H)** TNF. (**I**) Representative plots and quantification of the proportion of ABCs expressing IFNγ and TNF. Equivalent numbers of females and per group in all panels. (**A-C**) n=6-10 mice per group, data combined from two experiments. (**E-I**) n=6-8 mice per group, representative of two experiments. Each data point represents an individual mouse. Data presented as mean ± SEM. Analyzed by one-way ANOVA. P-values indicated as asterixis as follows: ****p<0.0001, *** p<0.001, ** p<0.01, * p<0.05.

To determine if the level of cytokine expression was distinct from other B cells, we compared the proportion of ABCs expressing IFNγ and TNF to non-ABC B cells. Specifically, we examined expression on ABCs compared to CD11c^-^Tbet^+^ and CD11c^-^Tbet^-^ B cells 35 days post-γHV68 infection (Figure 2D). A significantly increased proportion of ABCs expressed IFNγ and TNF compared to non-ABC B cells, both CD11c^-^Tbet^+^ and CD11c^-^Tbet^-^ B cells, in the spleen (Figure 2E). The mean fluorescent intensity (MFI) of IFNγ and TNF, of IFNγ or TNF positive cells, was significantly increased on ABCs compared to both non-ABC populations (Figure S2A, B). CD11c^-^ Tbet^+^ B cells expressed an intermediate level of IFNγ and TNF between ABCs and CD11c^-^Tbet^-^ B cells (Figure 2E, Figure S2A, B). Sex differences in ABC cytokine expression were not observed (Figure 2A-C, Figure S3 A-C). The high level of IFNγ and TNF cytokine expression indicates that ABCs may be functioning in a unique anti-viral capacity during latent infection.

To explore the relationship between latent γHV68 and the ABC population, we infected mice with ACRTA-γHV68, a recombinant strain of γHV68 in which the genes responsible for latency are deleted and a lytic gene, RTA, is constitutively expressed^30^. As a result, γHV68 infection is cleared following acute infection without ever establishing latency. We found that, at 35 days p.i., a time point at which ACRTA-γHV68 is cleared, ABCs are increased to the same level in the spleens of ACRTA-γHV68 infected mice as compared to spleens in mice infected with WT γHV68 (Figure 2F). Thus, ABCs persist in the absence of latent virus, indicating that ABCs act similarly to memory cells that remain following acute infection.

While the proportion of ABCs remained elevated in the absence of latent virus, cytokine expression was altered. The proportion of ABCs in ACRTA-γHV68-infected mice expressing IFNγ and TNF at 35 days p.i. was significantly reduced compared to those infected with WT γHV68 while remaining significantly elevated compared to naïve mice (Figure 2G-H). Additionally, the IFNγ^+^ ABCs in mice infected with ACRTA-γHV68 displayed a lower MFI than those from γHV68-infected mice, though the TNF MFI was not significantly different between the two groups (Figure S2C, D). Without the presence of the latent virus, we observe a decrease specifically in the proportion of ABCs that doubly express both IFNγ and TNF as well as an increase in ABCs that are negative for the expression of either IFNγ and TNF as compared to ABCs from WT γHV68-infected mice (Figure 2I). This finding indicates that ABCs respond to the latent virus by producing both IFNγ and TNF.

Together, these results show that ABCs are a predominant B cell subset expressing anti-viral cytokines IFNγ and TNF during latent γHV68. Further, our data establish that high levels of expression of these cytokines are dependent on the presence of the latent virus. These findings suggest that ABCs recognize the presence of the quiescent latent virus and respond by expressing anti-viral cytokines.

### ABCs are susceptible to γHV68 infection but are not a major viral reservoir

γHV68 is known to infect B cells, in particular germinal centre and memory B cells^31–34^. To determine if ABCs are directly infected by γHV68, we used a previously developed fluorescent strain of γHV68, γHV68.H2bYFP^35^ where fluorescence is easily detectable during acute infection, but over time falls off during latent infection. Mice were infected with γHV68.H2bYFP and 8 days p.i. spleens were collected, and flow cytometry was performed. We found that, during acute infection, a small proportion of ABCs were infected with γHV68. The same proportion of ABCs were positive for the fluorescent virus (1.2 ± 0.3%) as observed in previously activated B cells (1.1 ± 0.2%, Figure 3A-B), demonstrating that ABCs are not preferably targeted for infection. To our knowledge this is the first evidence of a virus directly infecting ABCs. We next asked what proportion of the γHV68 reservoir is made up of ABCs. The primary reservoir for γHV68 is previously activated B cells which aligns with our results that 73% of γHV68-infected cells are IgD^-^ B cells (Figure 3C). We found that ABCs make up only 6% of γHV68-infected cells (Figure 3C). Other infected cell populations included innate cells such as macrophages and DCs, in alignment with previous findings^31^, as well as a small proportion of NK and T cells (Figure 3C). We also examined the relationship between infection of ABCs and expression of IFNγ. We observed that during acute infection the proportion of ABCs expressing IFNγ was significantly lower in ABCs infected with γHV68 (4.3±1.3 %) as compared to uninfected ABCs (32.2±1.0%, Figure 3D). These results indicate that direct infection of ABCs was not driving IFNγ production and that ABCs are susceptible to γHV68-driven downregulation of IFNγ. These findings demonstrate that although a portion of ABCs are infected with γHV68, ABCs are not the major target for γHV68.

**Figure 3.**
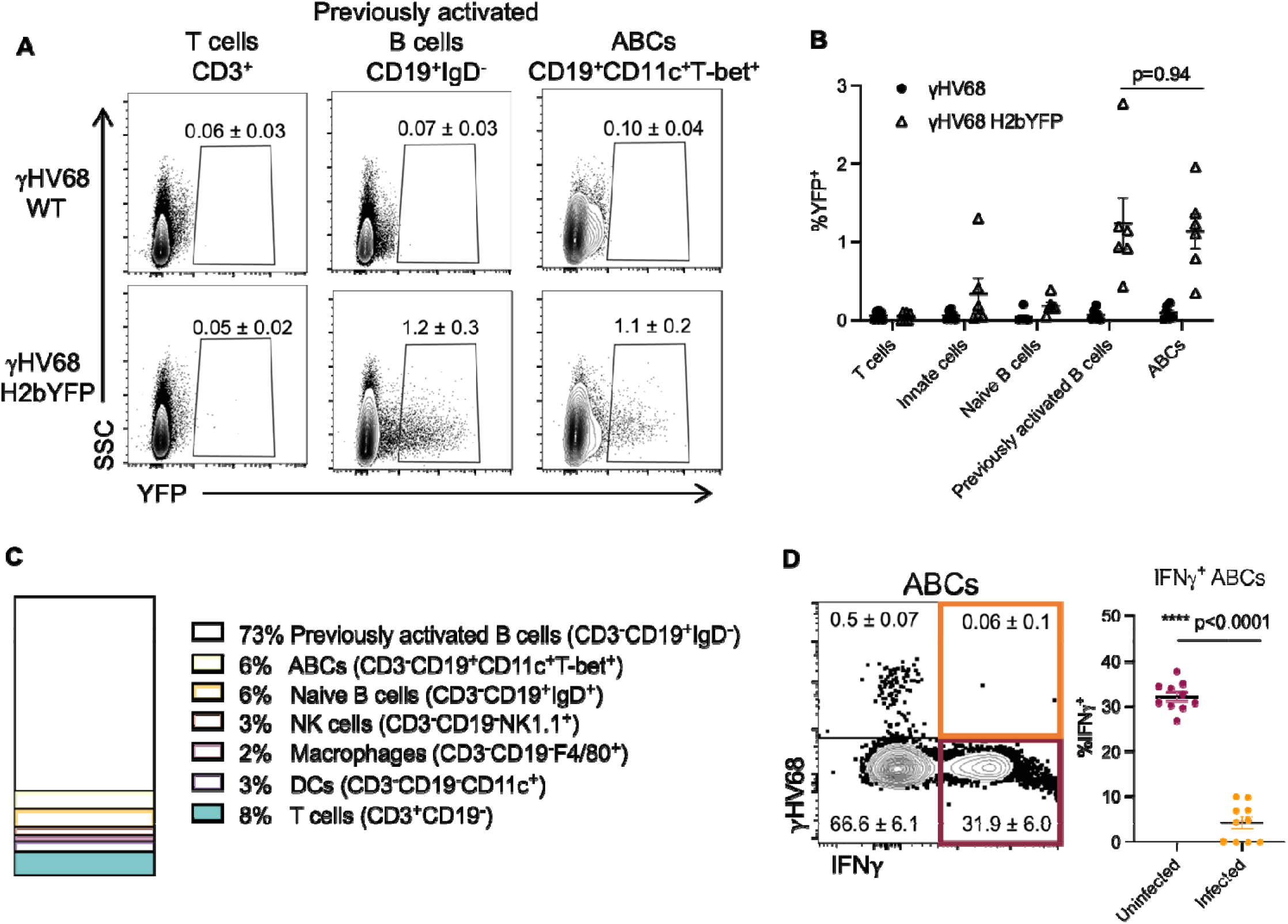
Analysis of ABC infection status with fluorescent γHV68 strain. Female C57BL/6(J) mice (6 to 8-week-old at infection) infected i.p. with γHV68 or γHV68.H2bYFP. 8 days p.i. spleen collected and processed for flow cytometry. (**A**) Representative flow cytometry plots displaying yellow fluorescent protein (YFP) expression (x-axis) versus side-scatter (SSC, y-axis) from mice infected with γHV68 or γHV68.H2bYFP with mean ± SEM. Samples gated on live, single lymphocytes, and then T cells (CD3^+^), total B cells (CD19^+^), previously activated B cells (CD19^+^IgD^-^), or ABCs (CD19^+^CD11c^+^T-bet^+^). Two samples per group concatenated for the purpose of visualization. Representative of two experiments. **(B)** Proportion of cell subsets positive for YFP expression from γHV68-infected mice (filled circles) and γHV68.H2bYFP-infected mice (open triangles). Each data point represents an individual mouse. (**C**) Proportion of YFP^+^ cells that are non-B cells (CD19^-^, white), non-ABC B cells (CD19^+^CD11c^-^T-bet^-^, grey), or ABCs (CD19^+^CD11c^+^T-bet^+^, black). Data n=6 mice per group, data compiled from two experiments, representative of three experiments. (**D**) Representative flow plot of ABCs (CD19^+^CD11c^+^T-bet^+^) in the spleen gated for YFP (γHV68.H2bYFP) and IFNγ with mean ± SEM. Dot plot displays the proportion of either uninfected (YFP^-^) or infected (YFP^+^) ABCs in the spleen that are positive for IFNγ. **(B, D)** Data presented as mean ± SEM, analyzed by Mann-Whitney test. Each data point represents an individual mouse. P-values indicated as asterixis as follows: ****p<0.0001.

### ABCs are dispensable for the control of acute infection and establishment of latency

To further examine the role(s) of ABCs in γHV68 infection, mice with a floxed B cell specific T-bet deletion were followed post-infection. Specifically, *Tbx21^fl/fl^Cd19^cre/+^* (KO) and littermate *Tbx21^fl/fl^Cd19^+/+^* (Ctrl) mice were infected with γHV68 for 6 or 35 days and the spleen was collected to examine the quantity of γHV68 by qPCR and the immune cell composition (Figure 4A). Mice were genotyped by PCR and loss of ABCs in KO mice was confirmed by flow cytometry (Figure 4B). No differences in clinical symptoms or weight changes were observed during γHV68 infection between Ctrl or KO mice (Figure S4A).

**Figure 4.**
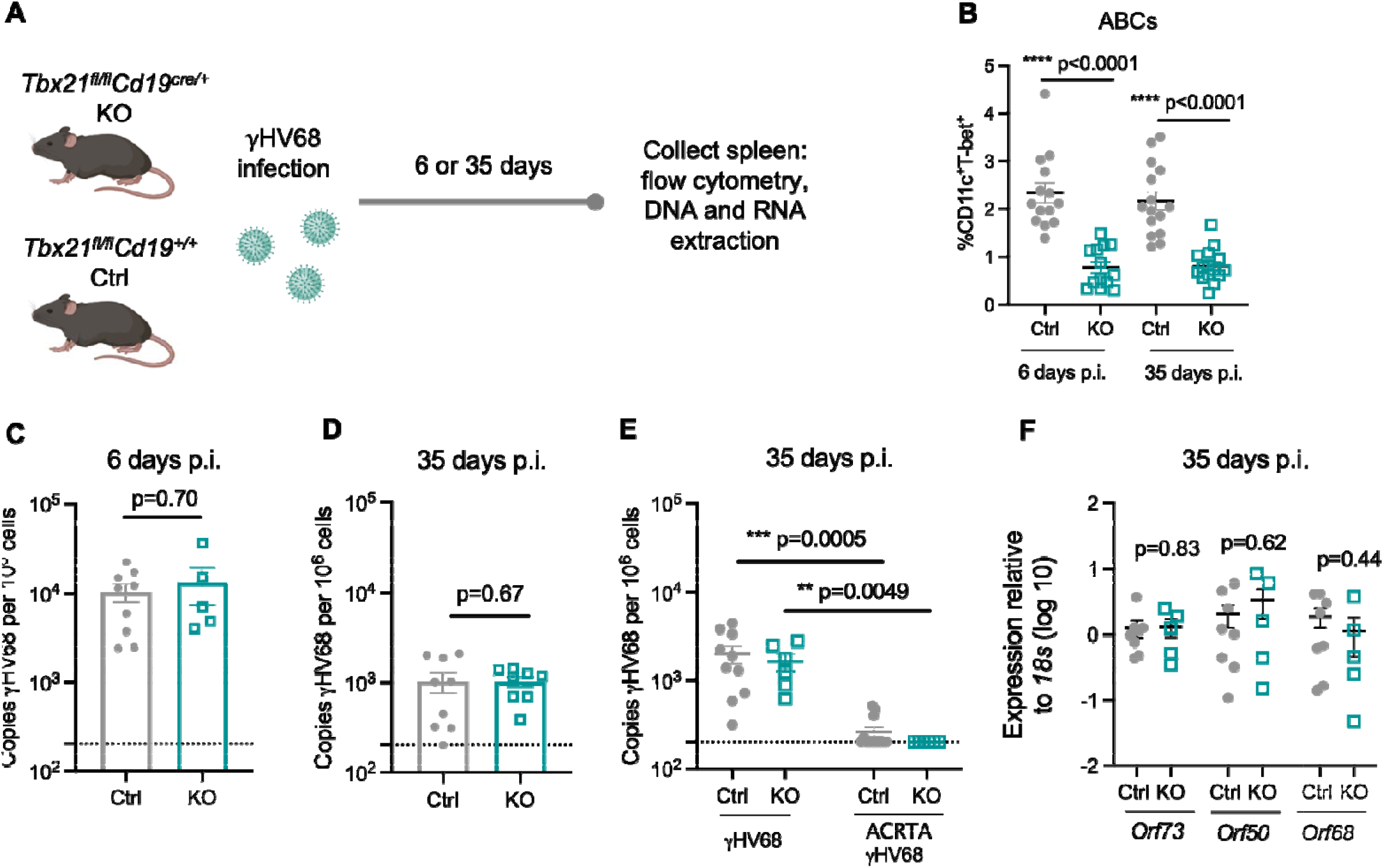
ABCs are not required for controlling lytic infection and achieving latency. *Tbx21^fl/fl^Cd19^+/+^* (Ctrl, filled grey circles) and *Tbx21^fl/fl^Cd19^cre/+^* (KO, open blue squares) mice were infected i.p. with γHV68 or ACRTA-γHV68. 6 or 35 days p.i., spleens collected and processed for RNA extraction, DNA extraction, and flow cytometry. (**A**) Experimental scheme for data shown in panels B, C, D, and F. (**B**) Proportion of ABCs (CD11c^+^T-bet^+^) of previously activated B cells (CD19^+^IgD^-^) in the spleen in Ctrl and KO mice 6 and 35 days post-γHV68 infection. (**C-D**) Splenic γHV68 viral load (copies *Orf50* per million cells) in Ctrl (filled circles) and KO (open squares) mice as determined by qPCR day 6 and 35 p.i., with limit of detection indicated by dotted line. (**E**) Splenic γHV68 viral load (copies *Orf50* per million cells) in Ctrl and KO mice infected with γHV68 or ACRTA-γHV68 35 days p.i.. (**F**) Relative expression in the spleen of *Orf73, Orf50*, and *Orf68* in Ctrl and KO mice at day 35 p.i.. Each data point represents an individual mouse. Data presented as mean ± SEM, analyzed by Mann-Whitney test (**B-D, F**) or Kruskal-Wallis H test (**E**).

To determine if knocking out ABCs alters immune cell composition, Ctrl and KO mice were infected or mock-infected with γHV68 and spleens analyzed at 6 or 35 days by flow cytometry to examine various T cell, B cell, and innate immune cell populations. No difference in the total number of splenocytes was observed (Figure S4B). The composition of the splenic immune profile was similar between Ctrl and KO mice with no significant differences observed in the relative proportions of B cells, CD8 T cells, DCs, neutrophils, NK cells, or macrophages between mock-infected mice or those infected with γHV68 for 6 or 35 days, though KO mice had greater proportions of CD4 T cells than Ctrl mice 35 days post-infection (Figure S5).

We then measured the quantity of γHV68 in the spleens of Ctrl and KO mice infected for 6 and 35 days by qPCR. We found no difference in that the quantity of γHV68 at 6 and 35 days p.i. between Ctrl and KO mice (Figure 4C-D) and the quantity of γHV68 did not differ between male and female mice (Figure S4C-D). To confirm that KO mice effectively control the lytic infection and develop latency, we infected Ctrl and KO mice with latency-deficient ACRTA-γHV68 for 35 days. We found that KO mice, like Ctrl mice, displayed similar viral loads that were near or below the limit of detection when infected with ACRTA-γHV68 (Figure 4E). As no virus was detected at day 35, mice lacking ABCs were effectively controlling the lytic infection. Furthermore, no differences were observed in the relative expression of viral genes associated with lytic infection (*Orf50, Orf68*) or latent infection (*Orf73*) between Ctrl and KO mice (Figure 4F). *Orf50* encodes the replication and transcription activator protein, which initiates viral lytic gene expression^36^. *Orf68* encodes a packaging protein that assists in moving the newly replicated viral genomes to the packaging motor, where they are loaded into capsids^37^. *Orf73* encodes for the latency-associated nuclear antigen that is required for the establishment and maintenance of latent infection^38,39^. As such, equal expression of lytic and latency-associated genes *Orf50, Orf68*, and *Orf73* between Ctrl and KO mice suggests that Ctrl and KO mice harbor similar levels of lytic and latent γHV68. Together, these results demonstrate that ABCs are not required for the clearance of acute infection and establishment of latency in steady-state conditions.

### Lack of ABCs results in a dysregulated γHV68 antibody response but does not alter the viral reservoir or anti-viral T cell response

ABCs are known to secrete anti-viral IgG2a/c^6^, and transfer of serum into mice without ABCs during LCMV infection was partially able to restore control of the infection^15^. To examine the antibody response in Ctrl versus KO mice, mice were infected with γHV68 for 35 days and sera was collected. The levels of anti-γHV68 IgG were the identical between Ctrl and KO mice, though Th1-associated IgG2c was significantly decreased in KO mice, while Th2-associated IgG1 was significantly elevated, when compared to Ctrl mice (Figure 5A-C). These results suggest that ABCs are the primary secreters of anti-γHV68 IgG2, though in their absence there is compensation by other B cell subsets resulting in increased anti-γHV68 IgG1 antibodies.

**Figure 5.**
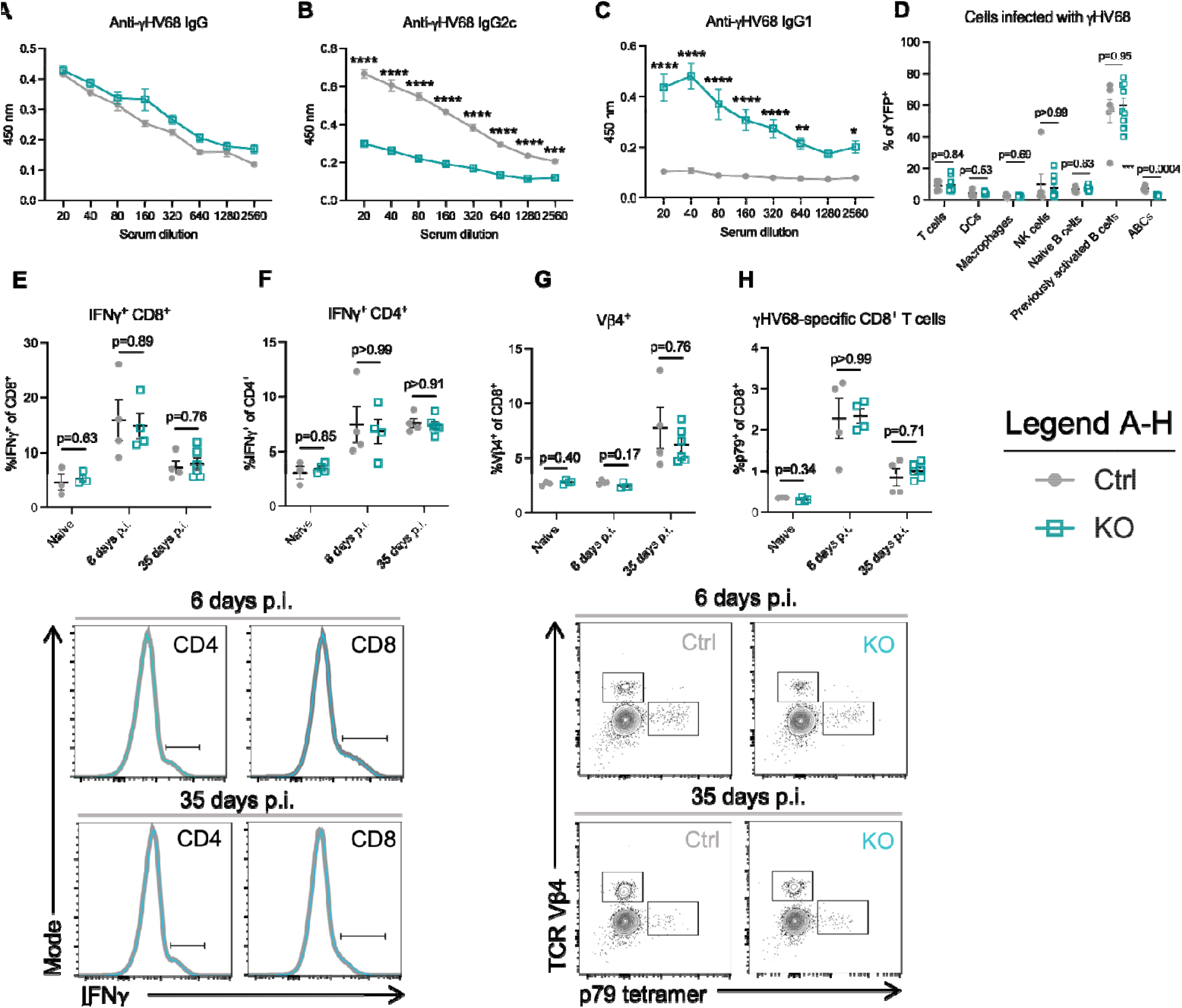
Mice without ABCs display a dysregulated anti-γHV68 antibody response, though no alterations to the cell populations infected with γHV68 or γHV68-responding T cells, compared to mice with ABCs. **(A-C)** *Tbx21^fl/fl^Cd19^+/+^* (Ctrl, filled grey circles) and *Tbx21^fl/fl^Cd19^cre/+^* (KO, open blue squares) mice were infected i.p. with γHV68 for 6 or 35 days. N=5 per group, representative of 2 experiments. (**D**) Ctrl and KO mice were infected i.p. with γHV68.H2bYFP for 8 days. N=6-8 per group, data compiled from 2 experiments. (**E-H**) Ctrl and KO mice were infected i.p. with γHV68 for 6 or 35 days, at which point spleens were collected for flow cytometry. Representative of 2 experiments. Each data point represents an individual mouse. Data presented as mean ± SEM. Analyzed by two-way ANOVA **(A-C)** or Mann-Whitney test **(D-H).** P-values indicated as asterixis as follows: ****p<0.0001, *** p<0.001, ** p<0.01, * p<0.05.

We next asked if the γHV68 reservoir was altered in mice without ABCs, as a portion of γHV68-infected cells were ABCs. In particular, we posited that an altered viral reservoir could impact the ability of KO mice to control γHV68 latency; previous findings indicate that various cell types infected by γHV68 may display differential susceptibility to reactivation^23,40^. To compare the viral reservoir in Ctrl and KO mice, we infected mice with a fluorescent strain, γHV68.H2bYFP, and, at 8 days p.i., examined immune cell populations that comprise the γHV68-infected population. We observed no difference in the proportion of infected cell populations, including T cells, DCs, macrophages, NK cells, and naïve and previously activated B cells (Figure 5D). This suggests that the cell populations infected with γHV68 are not substantially altered in KO mice compared to Ctrl mice, making it unlikely that changes to the reservoir impacted susceptibility to viral reactivation.

To ask whether ABCs influence anti-viral immune cells in the spleen, we next examined three cell populations previously shown to be important for control of latent γHV68: IFNγ-producing cells, γHV68-specific CD8^+^ T cells, and Vβ4^+^CD8^+^T cells. IFNγ is present at low levels during latency^41^ and is critical for controlling γHV68 reactivation from latency^23,24,42^. In particular, IFNγ-producing T cells are known to block γHV68 reactivation^23,43–45^. CD8^+^ T cells that harbor the Vβ4 TCR were also examined. CD8^+^Vβ4^+^ T cells expand following γHV68 infection and reach their highest levels during latency, wherein they persist throughout infection without taking on an exhausted phenotype^46–48^. γHV68-specific CD8^+^ T cells were also measured with tetramers to p79, an immunodominant γHV68 epitope for which CD8-specific memory T cells remain throughout latent infection^49^. We observed that the proportion of CD4^+^ and CD8^+^ T cells that express IFNγ were unchanged between mice with and without ABCs (Figure 5E-F). Further, no differences were observed in the proportion of either Vβ4^+^CD8^+^T cells or γHV68 p79-specific CD8^+^ T cells between mice with and without ABC circulating populations (Figure 5G-H). Additionally, we found that the proportion of NK cells and B cells that express IFNγ, and the MFI of IFNγ on these populations, was not changed between Ctrl and KO mice whether mock-infected or infected for 6 or 35 days (Figure S6). That a similar proportion of B cells expressed IFNγ indicates that another B cell subset likely compensates for the loss of IFNγ-expressing ABCs. Collectively, these findings indicate that ABCs are not acting to stimulate anti-viral immune cell populations in the spleen.

Together, these data indicate that knocking out ABCs leads to dysregulation of the γHV68 antibody response, without altering the γHV68 reservoir or T cell populations responding to the latent virus.

### Susceptibility to γHV68 reactivation increases in the absence of ABCs

We posited that the persistence of ABCs long-term during γHV68 and their secretion of IFNγ, TNF, and anti-viral antibodies suggest that ABCs have an important role in the suppression of γHV68 reactivation. To examine this role, ex vivo reactivation assays were performed on splenocytes from Ctrl and KO mice at 35 days p.i.. Splenocytes from γHV68-infected KO mice reactivated more frequently in culture than those from Ctrl mice (Figure 6A). This finding indicates that there is an increased propensity for γHV68 reactivation in the absence of ABCs, although the assay does not disentangle whether infected cells have an altered cell-intrinsic susceptibility to reactivation or if there is a cell-extrinsic influence on reactivation.

**Figure 6.**
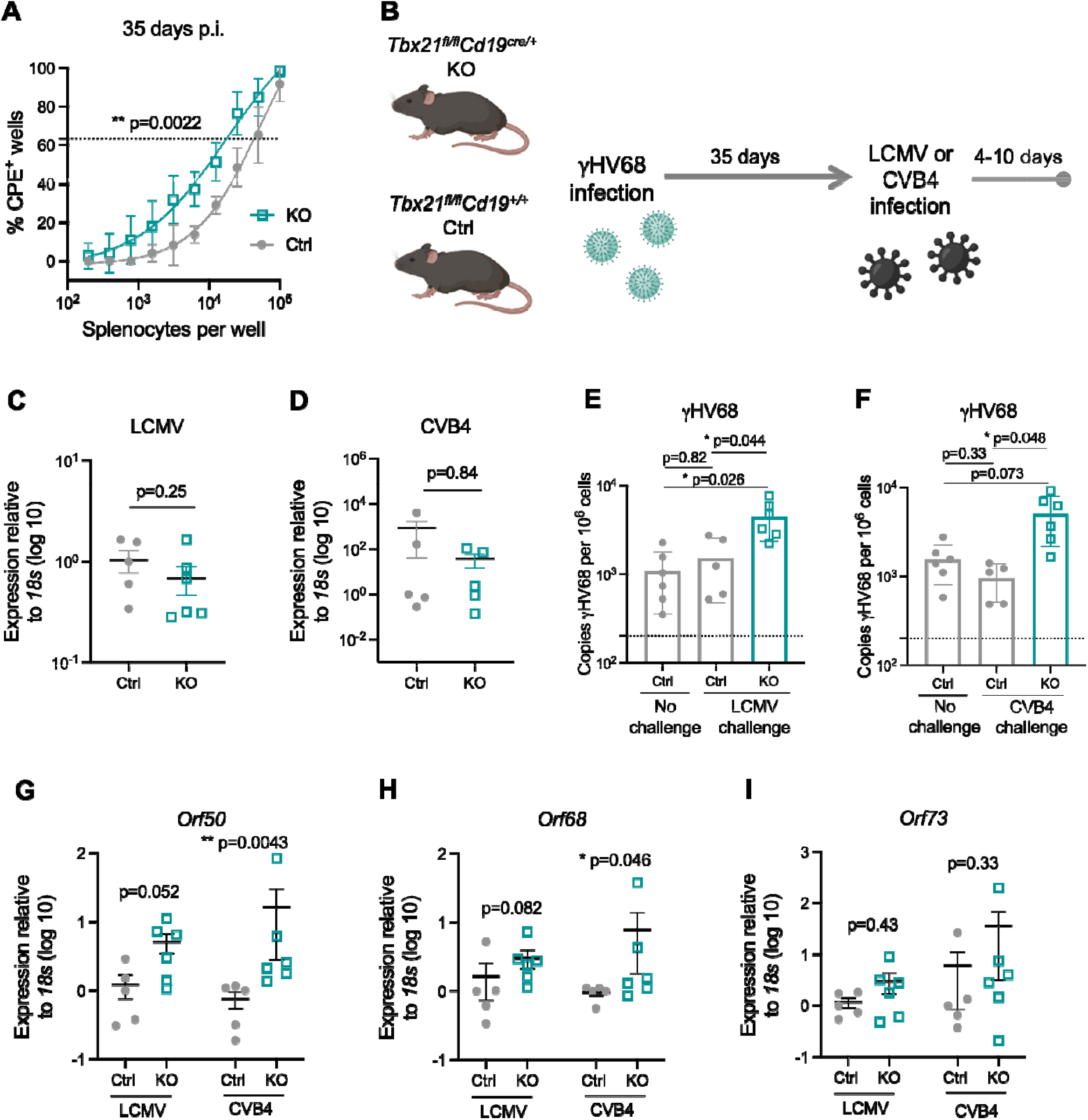
ABC knockout mice are more susceptible to reactivation. **(A)** *Tbx21^fl/fl^Cd19^+/+^* (Ctrl, filled grey circles) and *Tbx21^fl/fl^Cd19^cre/+^* (KO, open blue squares) mice were infected i.p. with γHV68. 35 days p.i., spleens were collected and processed for reactivation assay. N=6 mice per group, data compiled from two independent experiments. (**B**) Experimental scheme for data shown in panels C-I. Ctrl and KO mice were infected i.p. with γHV68 for 35 days and then challenged with either LCMV (Armstrong) or CVB4 for 10 to 4 days, respectively. (**C-I**) Spleens collected, processed, and DNA and RNA was extracted from total splenocytes following RBC lysis. **(C, D)** Relative expression of LCMV **(C)** and CVB4 **(D)** between Ctrl and KO mice measured by RT-qPCR. (**E, F**) Copies of γHV68 detected by qPCR in the spleen per million cells following challenge with LCMV or CVB4. (**G-I**) Relative expression in the spleen of *Orf73, Orf50*, and *Orf68* in Ctrl and KO mice. Each data point represents an individual mouse. Data presented as mean ± SEM, analyzed by Mann-Whitney test **(A, C-D, G-I)** or one-way ANOVA (**E, F**). P-values indicated as asterixis as follows: ** p<0.01, * p<0.05.

To further interrogate the role of ABCs in suppressing γHV68 reactivation, we next asked if mice without ABCs are more susceptible to γHV68 reactivation following infectious challenge. Heterologous viral infection was performed as a physiological mimic to reactivate latent γHV68. Importantly, we chose viruses that do not typically reactivate γHV68,^50^ LCMV and coxsackievirus B4 (CVB4). Specifically, KO and Ctrl mice were infected with γHV68 for 35 days and then challenged with LCMV or CVB4 (Figure 6B). No clinical symptoms were observed during either of the challenges in Ctrl or KO mice and there was no difference in the relative quantity of LCMV and CVB4 in the spleens of Ctrl versus KO mice (Figure 6C-D). Following challenge with LCMV or CVB4, mice lacking ABCs had elevated quantities of γHV68 in the spleen compared to mice with ABCs (Figure 6E-F). We reasoned that the increased γHV68 load following challenge may be due to increased reactivation and, in support, observed increased relative expression of two lytic-associated γHV68 genes, *Orf50* and *Orf68*, in KO mice compared to Ctrl mice following challenge with the heterologous viruses (Figure 6G-H). Alternatively, we did not observe a difference in expression of the latency-associated gene *Orf73*, between Ctrl and KO mice (Figure 6I). However, it remains possible that the increased quantity of γHV68 may result from expansion of infected B cells in response to the challenge, rather than reactivation. Collectively, these findings indicate that mice lacking ABCs may be more susceptible to γHV68 reactivation in the face of heterologous challenge.

## Discussion

Here we have shown that ABCs are increased during acute γHV68 infection and persist during latency for at least 150 days. Previous studies have shown that ABCs persist following clearance of viral infections^9^ and during chronic infection^15^, though their contribution during a latent infection was unexplored. Our results demonstrate a novel role for ABCs: they continuously respond in an effector manner to latent viral infection by secreting anti-viral cytokines and antibodies. Further, our results indicate that ABCs may play an important role in suppressing γHV68 reactivation during subsequent heterologous infectious challenges.

Our results show that ABCs continuously expresses anti-viral cytokines during latency and the presence of the latent virus is required for the sustained anti-viral cytokine expression by ABCs. Compared to other B cells, ABCs display uniquely high expression of anti-viral cytokines IFNγ and TNF during γHV68. In addition to expressing cytokines, ABCs also produce anti-viral antibodies, as deficiency of ABCs results in a significant loss of γHV68-specific IgG2c antibodies. We hypothesized that the production of anti-viral cytokines and antibodies by ABCs could be a way in which ABCs restrain γHV68 reactivation. Using mice deficient in ABCs, we showed that ABCs are important for the suppression of γHV68 reactivation in the face of heterologous viral challenge.

A surprising finding of our study is that ABCs are increased more so in female than male mice throughout γHV68 infection, in both the blood and spleen. Sex-differences are well-documented in anti-viral immune responses^51^ and ABCs have previously been shown to display a female sex bias in contexts of aging and autoimmunity^3,52^. It has recently been shown that the ABC female sex bias in lupus mice is abolished following the duplication of *Tlr7* in male mice^52^. While we show that there is no difference in the quantity of γHV68 between female and male mice, TLR7 stimulation is known to be important for ABC differentiation^2,3^ and TLR7 is critical for the control of lytic γHV68 infection and maintenance of latency^53^. While increased frequencies of ABCs were observed in female mice compared to males, we found that a similar proportion of the ABCs express anti-viral cytokines in both males and female mice. No differences between male and female mice were observed in viral load or reactivation.

Using a parallel approach to knockout T-bet in B cells with bone marrow chimeras, it was previously shown that mice lacking ABCs display an increased quantity of γHV68 at day 14 p.i. compared to controls^6^. This inconsistency with our results is most easily explained by a clear difference in the inoculating dose (lower herein) and the method of ABC knockout. Additionally, the two studies measure mice at different timepoints (day 14 p.i., the establishment of latency versus day 35 p.i., steady-state latency) and this likely hints at potential differences in the kinetics of latency establishment in mice lacking ABCs.

ABCs are known to possess an array of functional capacities and the precise mechanism(s) of ABC contribution to the restraint of γHV68 reactivation will be the focus of future studies. Here we have shown that only the antibody response is altered in mice deficient in ABCs and unexpectedly, changes are not observed in the anti-viral T cell response or viral reservoir. The ability of ABCs to continuously secrete anti-viral cytokines during latent infection is an additional mechanism that likely contributes to the suppression of γHV68 reactivation. Understanding the precise localization of ABCs in relation to γHV68-infected B cells will aid in our understanding of the development of splenic structures, in particular germinal centres, as ABCs have been shown to play a role in their development^54^.

Whether ABCs play a role in suppressing the reactivation of human gamma-herpesvirus infections such as Epstein-Barr virus (EBV) and Kaposi sarcoma-associated herpesvirus (KSHV), is not known. Understanding this role would have important relevance in diseases associated with these human gammaherpesviruses, including various malignancies, myalgic encephalomyelitis/chronic fatigue syndrome, and chronic-active EBV^55–57^.

This work demonstrates that ABCs are a long-lasting effector population during γHV68 latency and suggests that they are playing a unique role in the suppression of viral reactivation.

## Methods

### Mice

*Tbx21^fl/fl^Cd19^cre/+^* mice were generated by crossing *Tbx21^fl/fl^Cd19^cre/+^* and *Tbx21^fl/fl^Cd19^+/+^* mice. *Tbx21^fl/fl^* and *Cd19^cre/+^* mice were provided by Dr. Pippa Marrack^54^. C57BL/6(J) mice were originally purchased from The Jackson Laboratory. All animals were bred and maintained in the animal facility at the University of British Columbia. All animal work was performed per regulations of the Canadian Council for Animal Care (Protocols A17-0105, A17-0184).

### γHV68, ACRTA-γHV68, and γHV68H2B.YFP infection

γHV68 WUMS strain (ATCC), ACRTA-γHV68 (developed by Dr. Ting-Ting Wu, gift of Dr. Marcia A. Blackman)^30^, and γHV68H2B.YFP (developed and provided by Dr. Samuel H. Speck) were propagated in Baby Hamster Kidney cells (BHK, ATCC). Viruses were diluted in Minimum Essential Media (MEM) prior to infection and maintained on ice. 6- to 8-week-old mice were infected i.p. with 10^4^ PFU of γHV68, ACRTA-γHV68, γHV68H2B.YFP, or mock-infected with MEM. Clinical symptoms were not observed during γHV68, ACRTA-γHV68, or γHV68H2B.YFP infections in C57BL/6(J) nor *Tbx21^fl/fl^Cd19^cre/+^* or *Tbx21^fl/fl^Cd19^+/+^* mice.

### LCMV infection

LCMV Armstrong strain 53b (originally acquired from Dr. M.B. Oldstone) was propagated on BHK cells. Prior to infection, virus was diluted in RPMI-1640 media (Gibco) and maintained on ice. Mice (11- to 13-week-old) were infected i.p. with 2 × 10^5^ PFU LCMV or mock infected with media. No clinical symptoms were observed from LCMV infection in *Tbx21^fl/fl^Cd19^cre/+^* nor *Tbx21^fl/fl^Cd19^+/+^* mice.

### CVB4 infection

CVB4 Edward strain was propagated on HeLa cells and titred by plaque assay. Virus was diluted in Dulbecco’s Modified Eagle Medium (DMEM, Gibco) and maintained on ice prior to infection. Mice (11- to −13-week-old) were infected i.p. with 100 PFU CBV4. No clinical symptoms were observed from CVB4 infection in *Tbx21^fl/fl^Cd19^cre/+^* or *Tbx21^fl/fl^Cd19^+/+^* mice.

### Tissue harvesting and processing for flow cytometry

Mice were anesthetised with isoflurane and euthanized by cardiac puncture. For flow cytometry, blood was collected by cardiac puncture into 100 ul 0.5 M Ethylenediaminetetraacetic acid (EDTA) to prevent clotting and placed on ice until processing. Spleen extracted and placed into 2 ml PBS and kept on ice until processing. Blood was incubated in 10 ml warmed (37°C) ACK lysis buffer for 15 minutes at room temperature to remove red blood cells and washed twice with FACS buffer. Spleens were mashed through a 70 μm cell strainer with a 3 ml syringe insert to make a single cell suspension for each sample. Splenocytes were incubated in 4 ml warmed (37°C) ACK lysis buffer for 10 minutes on ice to lyse red blood cells and remaining cells were resuspended in FACS buffer and kept on ice until further use.

### Flow cytometry analysis of cell-type specific surface antigens and intracellular cytokines

For analysis of extracellular and intracellular antigens, 2 million cells per spleen sample or all cells collected per blood sample were stained with appropriate antibodies. Prior to staining, samples were incubated at 4°C covered from light for 30 minutes with 2 μl/ml Fixable Viability Dye eFluor506 (Thermo Fisher) while in PBS and then resuspended in rat anti-mouse CD16/32 (Fc block, BD Biosciences) antibody for 10 minutes. Then, samples were stained with fluorochrome labeled antibodies (Table 1) against cell surface antigens for 30 minutes covered from light at 4°C, washed, and resuspended in Fix/Perm buffer (Thermo Fisher) for 30 minutes-12 hours while maintained covered from light at 4°C. Samples were then washed twice with perm buffer and incubated 40 minutes with antibodies for intracellular antigens (Table 1) in Perm buffer at covered from light at RT. Cells were then washed and resuspended in FACS buffer with 2 mM EDTA. For analysis of cytokine production, 4 million cells per sample were stimulated ex vivo for 3 hours at 5% CO2 at 37°C in Minimum Essential Media (Gibco) containing 10% fetal bovine serum (FBS, Sigma-Aldrich), 1 μl/ml GolgiPlug (BD Biosciences), 10 ng/ml PMA (Sigma-Aldrich) and 500 ng/ml ionomycin (Thermo Fisher) and washed prior to staining. Samples were collected on an Attune NxT Flow Cytometer (Thermo Fisher) and analyzed with FlowJo software v10 (FlowJo LLC).

**Table 1.**
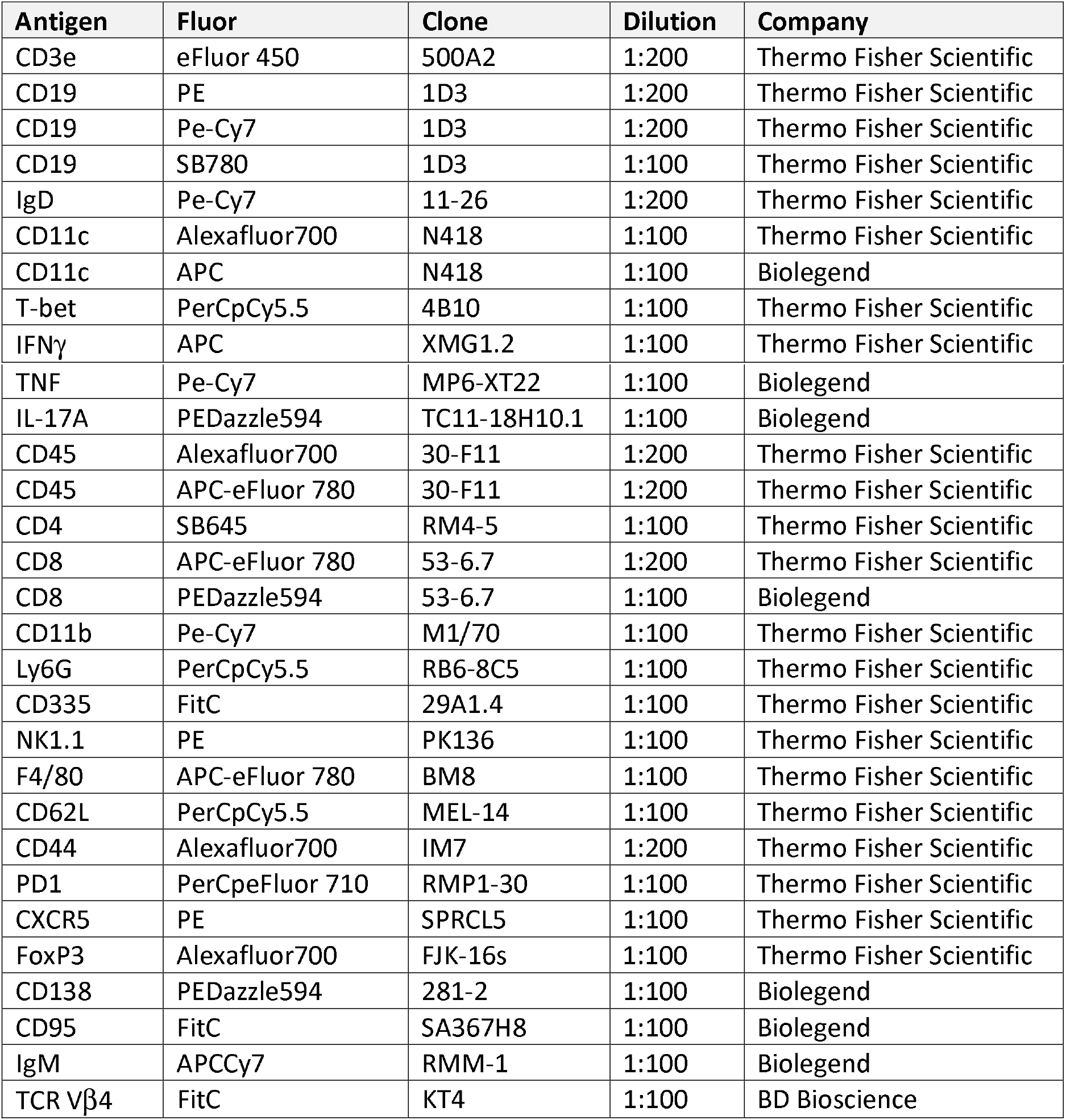
Flow cytometry antibodies

### Tetramer staining

γHV68-specific CD8 T cells were identified by staining with a tetramer acquired from the NIH Tetramer Facility. 2 million cells per well were incubated for 1 hour at RT with p79 (K^b^/ORF61_524–531_ TSINFVKI, diluted 1:400) following viability staining and Fc receptor block, as described above, but before extracellular antigen staining. Uninfected mice stained with tetramers used as negative gating controls.

### γHV68 qPCR

DNA was isolated from 4 × 10^6^ splenocytes using PureLink™ Genomic DNA Mini Kit (Thermo Fisher) and quantified with a spectrophotometer. qPCR was performed using 2x QuantiNova Probe Mastermix (Qiagen, USA) on the Bio-Rad CFX96 Touch™ Real Time PCR Detection system. Copies of γHV68 were quantified in duplicate wells with 150 ng DNA per reaction using primers and probes specific to γHV68 *Orf50* and mouse *Ptger2* (Table 2). Standard curves were obtained by serial dilutions of *Orf50* and *Ptger2* gBlocks (*Orf50*: 2×10^6^ – 2×10^1^; *Ptger2*: 5×10^7^-5×10^2^) that were amplified in parallel.

**Table 2.**
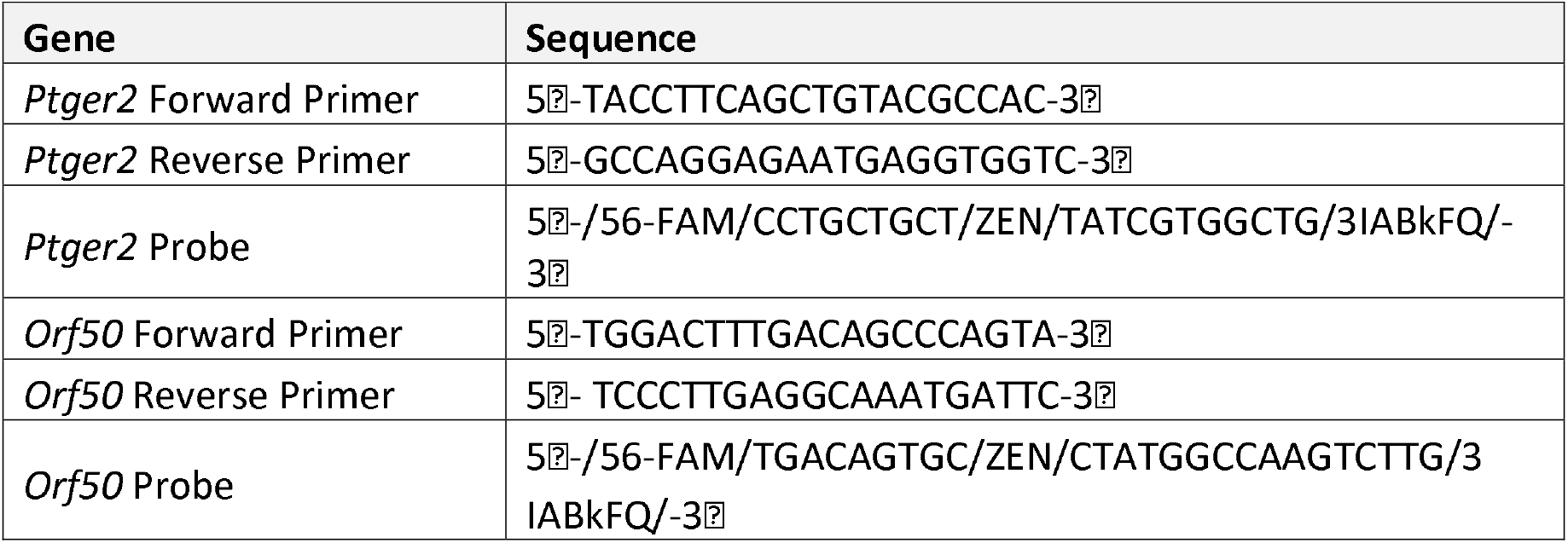
γHV68 qPCR primers and probe sequences

### RT-qPCR for γHV68 lytic and latency-associated genes and relative quantity of LCMV and CVB4

Portion of spleen stabilized in RNAlater immediately following collection and stored at −80°C for up to 12 weeks. RNA extracted with RNeasy mini kit (Qiagen, cat no. 74104) and cDNA immediately synthesized with High-Capacity cDNA Reverse Transcription Kit (Thermo Fisher, cat no. 4368814). Reaction performed with 500ng per reaction in duplicate wells. cDNA quantified with a spectrophotometer. qPCR was performed using iQTM SYBR^®^ Green supermix (Bio-Rad) on the Bio-Rad CFX96 Touch™ Real Time PCR Detection system. Transcript specific primers have been previously described for *Orf50, Orf73*, and *Orf68*^58^, LCMV^59^, and CVB4^60^. Primers were ordered from Integrated DNA Technologies (Table 3). Normalized to the ribosomal housekeeping gene *18s* and expression determined relative to control group.

**Table 3.**
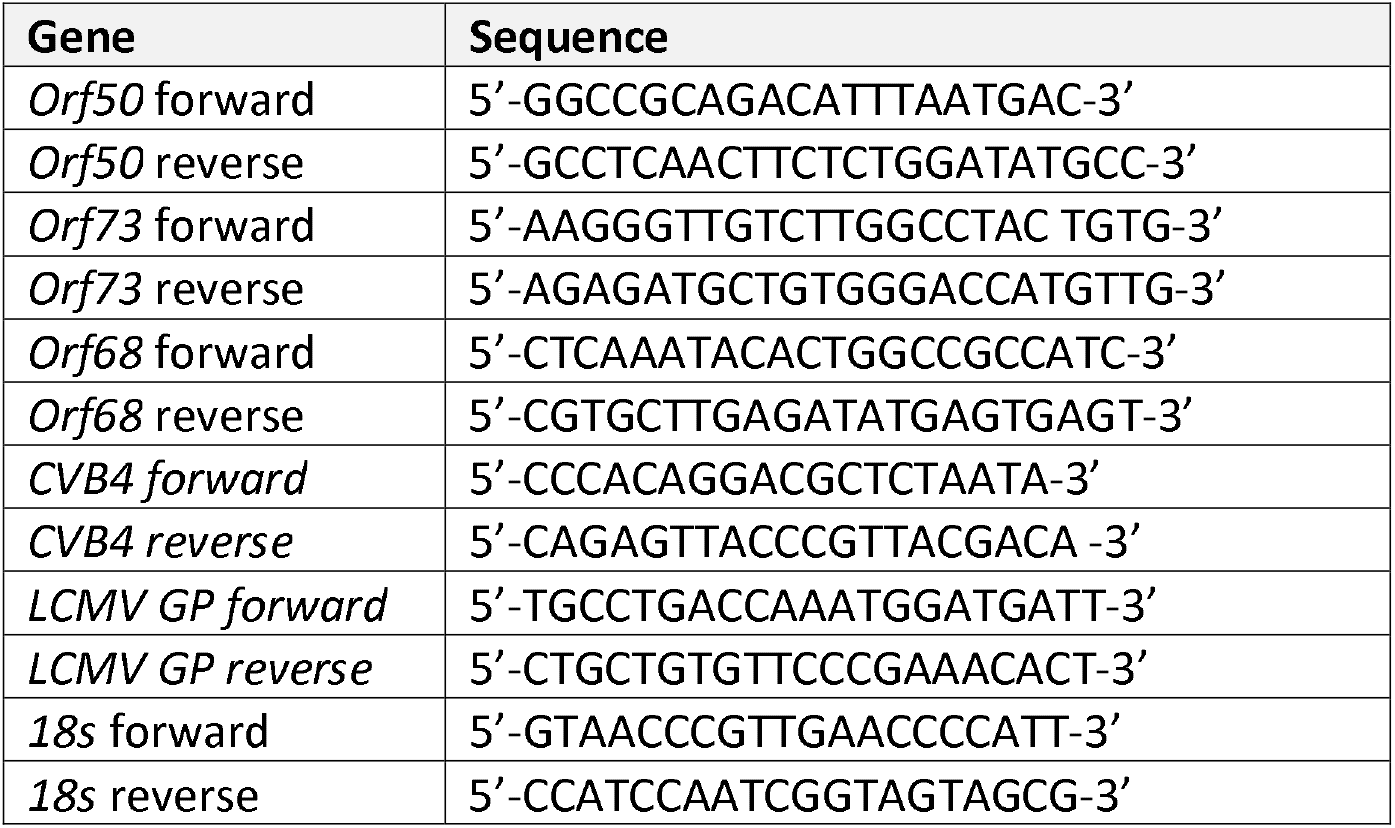
qPCR for lytic and latent γHV68 genes primer sequences

### Ex vivo limiting dilution reactivation assay

A single cell suspension following RBC lysis with ACK lysis buffer was prepared from freshly collected spleens. To analyze ex vivo reactivation, limiting dilution reactivation assay was performed as previously described^17^. Two-fold serial dilutions of splenocytes starting at 10^5^ cells per well were plated onto monolayer of mouse embryonic fibroblast (MEF, C57BL/6) cells (ATCC) in 96-well flat-bottom polystyrene tissue culture plates. Twelve wells were plated per dilution. Plates were incubated at 37°C 5% CO_2_ for 10 days and scored for microscopically for viral cytopathic effects (CPE). Data were plotted as sigmoidal dose curves and interpolation was used to determine the cell density per sample at which 63.2% of wells were positive for CPE.

### Anti-γHV68 antibodies

Serum was collected via cardiac puncture and maintained at RT to allow for clotting. The sera were isolated by centrifugation 2000 × g for 10 min, aliquoted, and stored for up to 12 months at −80°C prior to running the ELISA. Anti-γHV68 antibodies were quantified by standard indirect ELISA. Briefly, γHV68 virions were inactivated in 4% paraformaldehyde (PFA) for 20 minutes at RT. Then, ELISA plates (NUNC, Thermo Fisher) were coated with 5 μg/ml γHV68 in coating buffer (0.05 M NaHCO_3_, pH 9.6) with 1% PFA overnight at 4°C. The plate was then washed 4x with wash buffer (PBS-0.05% Tween-20), blocked in blocking buffer (5% NBCS in PBS) for 1 hour at 37°C, and incubated with serial dilutions (1:20, 1:40, 1:80, 1:160, 1:320, 1:640, 1:1280, 1:2560) of test sera diluted in blocking buffer for 2 hours at 37°C. The plate was washed 4x with wash buffer and bound antibody was incubated with HRP-conjugated goat anti-mouse IgG (Thermo Fisher), rat anti-mouse IgG1 (BD Biosciences), or goat anti-mouse IgG2c (Thermo Fisher), all diluted 1:500 in blocking buffer, for 1 hour at 37°C, washed 4x with wash buffer, and detected by TMB substrate (BD Biosciences). Absorbance was read at 450 nm on a VarioSkan Plate Reader (Thermo Fisher).

### Statistical analyses

Data analysis and presentation as well as statistical tests were performed using GraphPad Prism software 8.4.2 (GraphPad Software Inc.). Results are presented as mean ± SEM. Statistical tests, significance (p-value), sample size (n, number of mice per group) and number of experimental replicates are stated in the figure legends. Statistical analyses included: two-way ANOVA with Geisser-Greenhouse’s correction, Mann-Whitney test, and one-way ANOVA. P-values indicated by asterisks as follows: ****p<0.0001, *** p<0.001, ** p<0.01, * p<0.05.

### Study approval

All work was approved by the Animal Care Committee (ACC) of the University of British Columbia (Protocols A17-0105, A17-0184).

## Acknowledgements

The following reagent was obtained through the NIH Tetramer Core Facility: class I p79 tetramer. The authors thank Martin Richer for the helpful feedback on the manuscript.

