## Supplemental material for "Age-associated B cells are long-lasting effectors that restrain reactivation of latent γHV68"

**Figure S1**

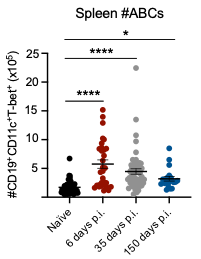

**Figure S1: Total number of ABCs are increased during γHV68 infection**

C57BL/6(J) mice (6–8-week-old at infection) were mock-infected with MEM (naïve) or infected i.p. with γHV68 for 6, 35, or 150 days. Spleens were then collected and processed for flow cytometry. Data compiled from multiple experiments, presented as mean ± SEM, analyzed by one-way ANOVA, P-values indicated as asterixis as follows: ****p<0.0001, * p<0.05.

**Figure S2**

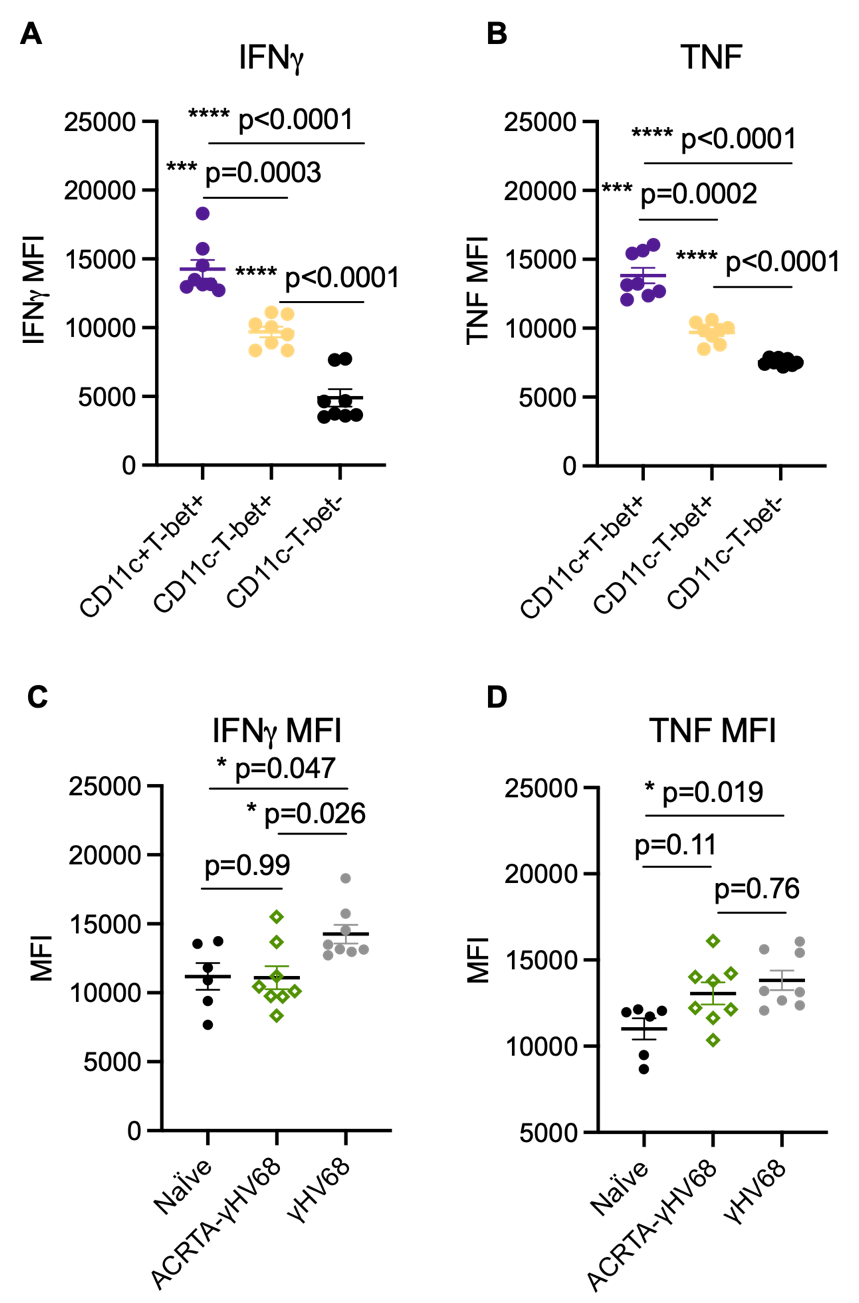

**Figure S2: IFNγ and TNF MFI on ABCs and non-ABCs during γHV68 infection**

(**A, B**) C57BL/6(J) mice were infected i.p. with γHV68. 35 days p.i. the mean fluorescence intensity (MFI) of IFNγ and TNF was examined on three CD19^+^ B cell populations in the spleen: CD11c^+^T-bet^+^ (purple), CD11c^-^T-bet^+^ (yellow), and CD11c^-^T-bet^-^ (black). (**C, D**) C57BL/6(J) mice were infected i.p. with ACRTA-γHV68 (green), WT γHV68 (grey), or mock-infected with MEM (black) for 35 days. Spleens were collected and the MFI of IFNγ^+^ and TNF^+^ ABCs (CD19^+^CD11c^+^T-bet^+^) was examined. Data presented as mean ± SEM, analyzed by one-way ANOVA, * p<0.05.

**Figure S3**

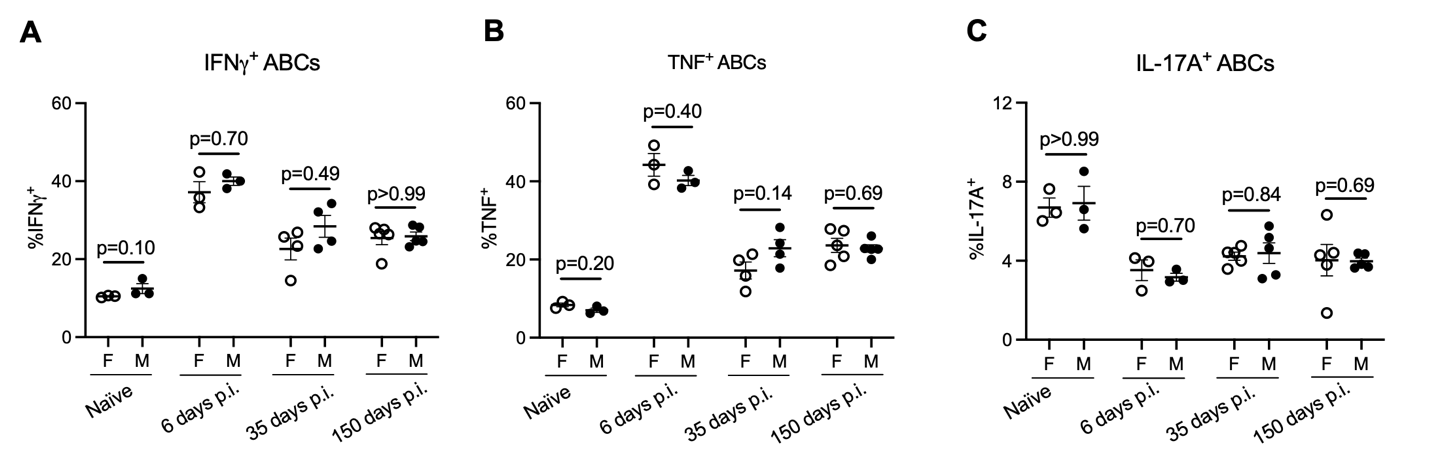

**Figure S3: Cytokine production by ABCs does not vary based on sex.**

C57BL/6(J) mice (6–8-week-old at infection) were mock-infected with MEM (naïve) or infected i.p. with γHV68 for 6, 35, or 150 days. Spleens were then collected and processed for flow cytometry. Proportion of ABCs in the spleen expressing (**A**) IFNγ, (**B**) TNF, or (**C**) IL-17A. Same data as Figure 4.2, separated by females (F) and males (M). Representative of two independent experiments. Data presented as mean ± SEM. Analyzed by Mann-Whitney test.

**Figure S4**

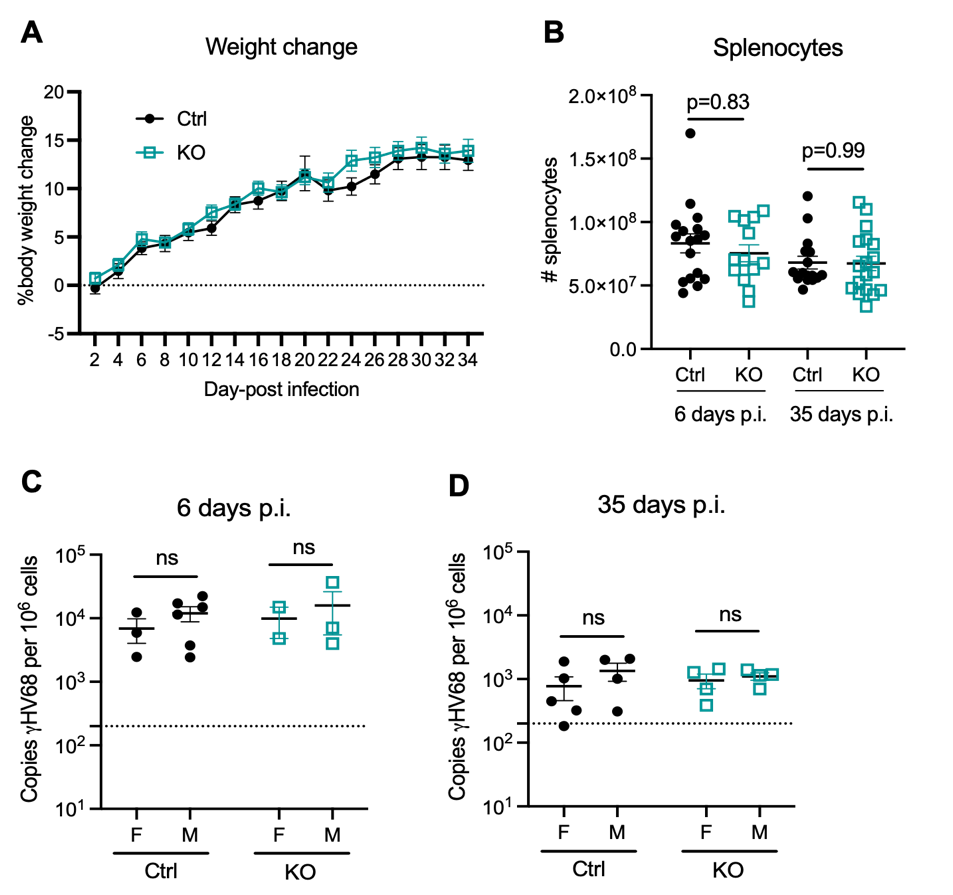

**Figure S4: Mouse weight change during γHV68, splenocyte number upon collection, and viral load by sex.**

*Tbx21^fl/fl^Cd19^+/+^* (Ctrl, filled grey circles) and *Tbx21^fl/fl^Cd19^cre/+^* (KO, open blue squares) mice were infected i.p. with γHV68. (**A**) Ctrl and KO mice were weighed every two days over the course of γHV68 infection, and the percentage body weight change from the day of infection was calculated. N=14-17 mice per group, data combined from three independent experiments. (**B**) The number of live splenocytes from mice infected with γHV68 for 6 or 35 days were counted by hemacytometer. Data combined from five independent experiments. (**C, D**) *Tbx21^fl/fl^Cd19^+/+^* (Ctrl, filled grey circles) and *Tbx21^fl/fl^Cd19^cre/+^* (KO, open blue squares) mice were infected i.p. with γHV68 for 6 or 35 days p.i.. Quantity of γHV68 in the spleen was measured by qPCR in female (F) and male (M) mice. Data presented as mean ± SEM. Analyzed by Mann-Whitney test.

**Figure S5**

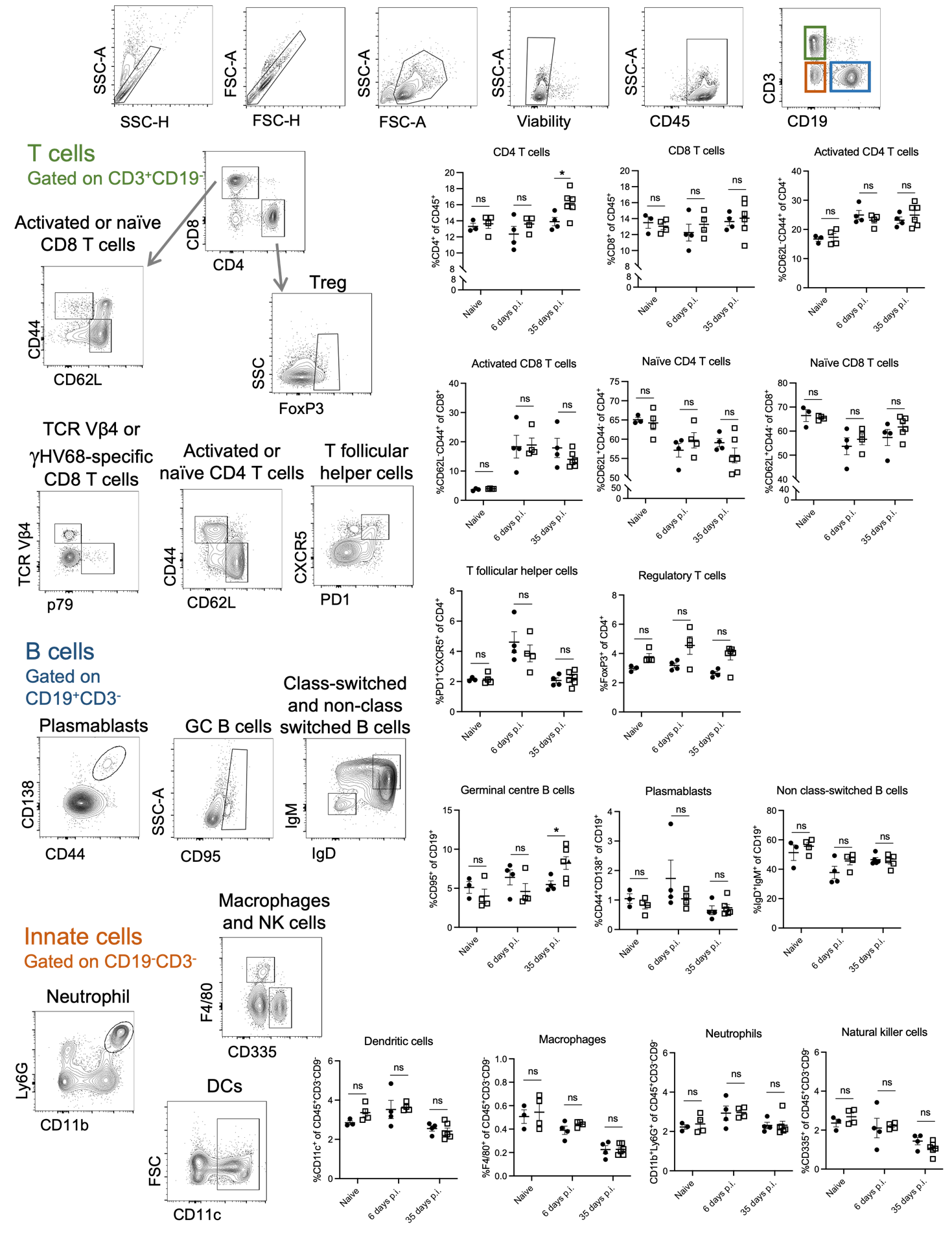

**Figure S5: Immune cell subsets in Ctrl versus KO mice during γHV68 infection.**

*Tbx21^fl/fl^Cd19^+/+^* (Ctrl, filled circles) and *Tbx21^fl/fl^Cd19^cre/+^* (KO, open squares) mice were mock infected (naïve) or infected i.p. with γHV68 for 6 or 35 days. Immune cells in the spleen examined by flow cytometry, including CD19^+^ B cells, CD4^+^ T cells, CD8^+^ T cells, CD11c^+^ dendritic cells, CD11b^+^Ly6G^+^ neutrophils, CD335^+^ natural killer cells, and F4/80^+^ macrophages. Representative gating scheme and proportion of cell subsets shown. Data representative of 2 experiments. Data presented as mean ± SEM. Analyzed by one-way ANOVA, * p<0.05.

**Figure S6**

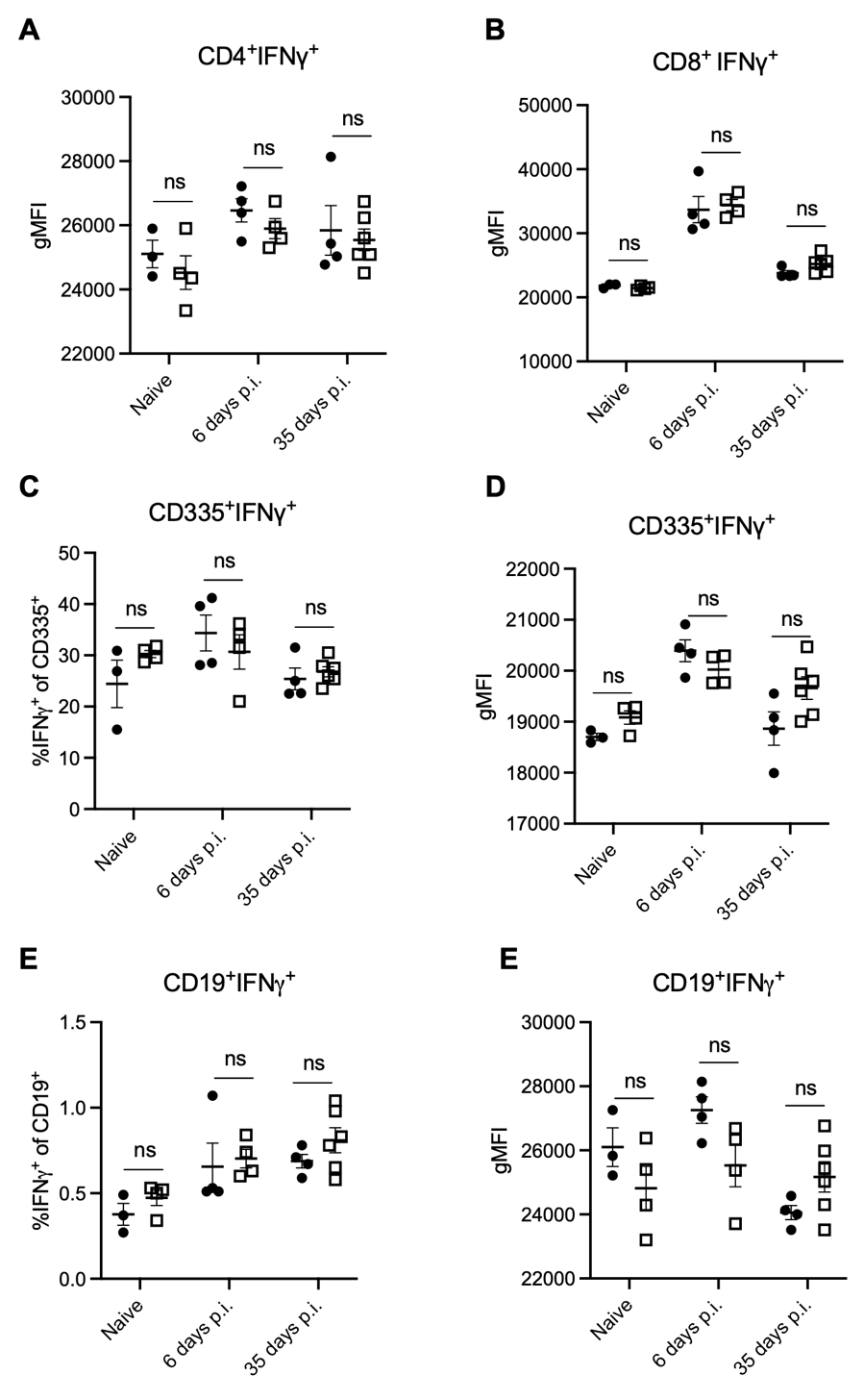

**Figure S6: No major changes to cells known to play a role in the control of γHV68 latency in KO mice**

*Tbx21^fl/fl^Cd19^+/+^* (Ctrl, filled circles) and *Tbx21^fl/fl^Cd19^cre/+^* (KO, open squares) mice were infected i.p. with γHV68. 6 or 35 days p.i., spleens were collected and processed for flow cytometry. Examination of the proportion of CD4 T cells (A) and CD8 T cells (B) expressing IFNγ and IFNγ gMFI in CD4 (C) and CD8 (D) T cells. Examination of the proportion of NK cells (E) and B cells (F) expressing IFNγ and IFNγ gMFI in NK cells (G) and B cells (H). Proportion of Vβ4^+^of CD8^+^ T cells (I). Proportion of CD8 T cells specific for γHV68 p79 antigen (J). Data presented as mean ± SEM, analyzed by one-way ANOVA.
